# Bayesian Network Analysis Identifies PREX1 as a Master Regulator of Cytoskeletal Disruption in Inflammatory Bowel Disease

**DOI:** 10.64898/2026.07.24.740476

**Authors:** Snezhanna Medvedeva, Julia Kulygina, Ksenia Morozova, Julia Popova, Maksim Struchalin, Marina Osipenko, Elena Kozhevnikova

## Abstract

**Background:** Inflammatory bowel diseases are complex multifactorial and polygenic conditions with incompletely understood etiology. Transcriptomic and metabolomic studies reveal hundreds of genes and metabolites associated with the inflammatory dowel diseases (IBD), while about 240 genetic loci are associated with these diseases. IBD are accompanied by numerous epithelial lesions including epithelial cells damage and barrier dysfunction. Recent studies suggest that cytoskeleton dysregulation underlies these aspects of IBD. We have previously shown that the disruption of cortical actin filaments and microvilli degeneration are characteristic of *Muc2* mouse IBD model.

**Methods:** We used transmission electron microscopy to evaluate ultrastructural defects in the intestinal epithelium of IBD patients. A probabilistic graphical model (Bayesian network) was employed in order to identify the key gene network hierarchy that might define cytoskeletal architecture among other regulatory events in IBD. The Bayesian network was based on preselected key node genes from open access transcriptomic data obtained in patients with Crohn’s disease.

**Results:** Electron microscopy of IBD patients demonstrated disruption of the brush border ultrastructure, similar to the microvillar defects previously described in *Muc2* colitis model mice. We highlighted a list of genes contributing to actin polymerization and bundling, of which *PREX1* were further confirmed using the transcriptome data obtained from *Muc2* mice.

**Conclusion:** Our study underscores the utility of Bayesian network analysis in complex cellular phenotypes that helped to identify potential target genes responsible for cortical cytoskeleton and microvilli disruption upon chronic intestinal inflammation.

## 1. Introduction

Inflammatory bowel diseases (IBD) represent a significant growing health challenge all over the world. These disorders include ulcerative colitis (UC), Crohn’s disease (CD) and indeterminate colitis, all of which are characterized by chronic, persistent and relapsing inflammation of the gastrointestinal tract. IBD is described as a complex multifactorial disorder that arises from an interplay between environmental factors (including internal environment such as metabolome and microbiota), genetic background and epigenetic regulation (1).

One of the key features of IBD is epithelial damage, which manifests at both the tissue and cellular levels. Tissue injury includes ulcerations and cell death upon chronic inflammation, while cell-level damage involves ultrastructural changes in mitochondria, disruption of intracellular junctions, microvilli loss, disorganization of terminal web and cytoskeletal structure (2–5). The same phenotypic manifestations characterize animal models of IBD (6,7). These aspects are being actively investigated using highthroughput sequencing data in order to identify key regulatory genes involved in structural damage of epithelium (8).

It is proven that relatives of CD-affected individuals have increased IBD genetic risk (9). Largescale genome-wide association studies and other genomic analyses have established more than 240 candidate loci for IBD (10). They include genes involved in epithelial barrier function (*CDH1, CLDN4, MUC3A, MYO9B*), innate and adaptive immune response (*NOD2*, *HLA* genes, *IL10R*, *IL23R*, *TNFSF15*, *PTPN22*) and autophagy (*ATG16L1*, *ULK1*, *IRGM*), emphasizing the disturbance of intestinal homeostasis and exaggerated host-microbiota immune activity in IBD (1,11,12). Most of these genes are of high IBD risk, but they explain only a small portion of the variance in susceptibility to UC and CD (1). Since IBD is a complex polygenic condition, the cumulative effects of multiple genetic factors play a more significant role in the overall disease susceptibility than the presence of individual risk alleles. Transcriptomic analysis has been widely used to discover novel target genes and understanding pathogenesis of IBD. However, the typical number of genes discovered using differential gene expression reaches hundreds or thousands (13,14). This might be challenging to identify the most relevant gene targets given the number of differentially expressed genes (DEGs).

To advance drug discovery and elucidate the genetic architecture of IBD, it is essential to prioritize candidate genes based on their functional relevance. The most widely used method for this purpose is Weighted Gene Coexpression Network Analysis (WGCNA) (15). WGCNA estimates pairwise correlation among gene transcripts and uses this information to estimate interconnections (co-expression) between genes groping them into co-expression modules (16). The ones that are highly connected with other genes in the module are considered as hub genes. These module representative genes are often considered as the most influential players within transcription network.

These hub genes can be used further to evaluate their conditional interactions and predict gene regulation hierarchy with a Bayesian network (17). A Bayesian network (BN) is a probabilistic graphical model which represents relationships between variables using a directed acyclic graph (18). BNs have been used in studying gene expressions to reveal causal relationships among genes and transcriptional gene regulatory networks (19–22). Thus, hub genes evaluated in WGCNA are used as nodes, while edges represent associations between them and the direction of edges shows causality. Such approach was successfully used to classify hematological malignancies (17).

In this study, we employed transmission electron microscopy to demonstrate that ultrastructural alterations in microvilli architecture and actin cytoskeleton organization are remarkably similar in the intestinal epithelium of both IBD patients and *Muc2*-deficient mice, an established model of IBD. We further elucidated a gene regulatory network by applying the WGCNA method, followed by Bayesian network analysis, to open-access gene expression data derived from IBD patients. This approach identified genes with the highest impact on cytoskeleton and epithelial integrity. A transcriptomic analysis of gene expression in colon of *Muc2* knock-out (KO) mice revealed *PREX1* as a universal cytoskeletal regulator in the intestinal epithelial cells (6).

## 2. Materials and Methods

### 2.1. Data sources

This study utilized transcriptomic, metabolomic, and clinical data obtained from the published research by Braun et al.0 (23). The original study included two cohorts: Chinese and Israeli; however, in this work, we used the Israeli cohort to build the Bayesian Network, which was tested as calprotectin predictive model on the Chinese cohort. The Israeli cohort from the source study included 45 participants: 17 newly diagnosed Crohn’s disease (CD) patients and 25 healthy controls. Age distribution was similar for CD patients and controls. The Chinese cohort included 20 healthy individuals and 20 CD. Chinese cohort original data does not include calprotectin in metadata.

Gene expression data were downloaded from the GSE199906 and GSE233900 datasets available in the Gene Expression Omnibus (24). Clinical data were extracted from the supplementary materials of the study by Braun et al. (23). Clinical data included parameters such as age, sex, disease status, C-reactive protein (CRP) levels, and calprotectin levels (for Israeli data only).

### 2.2. Differential expression, WCGNA, and gene ontology analysis

Differential expression analysis of patients’ genes was performed using the edgeR package (version 4.4.1) (25). Differential gene expression between the control and patient groups was performed using quasi-likelihood F-tests and adjusted for age and sex. For genes, the thresholds for differential expression were set at log_2_ of fold change > 1 and FDR < 0.05. DEGs were further analyzed to using WCGNA package (15) to predict co-expression gene modules. Intra-modular connectivity was computed for each gene to determine its relative importance within its assigned module. For hub gene identification, we integrated three complementary metrics: intra-modular connectivity, module membership, and aggregate gene-trait associations. These three metrics were standardized via Z-score normalization and combined into a composite hub score using equal weighting. The top 20 genes per module were selected as hub candidates based on this integrated score. From the selected genes, those encoding immunoglobulin chains were omitted from further analysis.

### 2.3. Construction and analysis of the Bayesian Network

The BN was constructed using the bnlearn R package (version 5.0.2) for the hub genes (26). The BN structure was inferred using the hill-climbing algorithm with the Bayesian Information Criterion for Gaussian data. To ensure robustness, bootstrap resampling (100 iterations) was performed, and an averaged network was derived from arcs with consistent directional support (strength > 0.7). The final network retained only high-confidence edges (bootstrap strength ≥ 0.7 and direction probability > 0.7). Networks were visualized using Rgraphviz (version 2.50.0) (27) and igraph (version 2.1.4) (28) packages, with nodes colored by their WGCNA module affiliation. Node size was scaled by out-degree centrality to highlight topological influence. In figures, arrows, nodes and captions were slightly adjusted using Adobe Illustrator 2022 for better visibility.

Genes participating in high-confidence edges (strength ≥ 0.7) were subjected to GO enrichment analysis using clusterProfiler (version 4.14.6) (29). Human Entrez IDs were mapped to gene symbols via org.Hs.eg.db (version 3.20.0) (30). Enriched terms in the Cellular Component ontology were identified with thresholds of *p* < 0.05 (adjusted using Benjamini*-*Hochberg’s method). Results were visualized via dot plots and exported for further interpretation. For calprotectin prediction, the BN was constructed using calprotectin as a network node, while outgoing connections from calprotectin were blacklisted. Predictions of calprotectin levels were generated using Bayesian likelihood weighting. For the Israeli dataset, predicted calprotectin was plotted against its actual data. For the Chinese dataset, no information on calprotectin was provided, thus, we used BN-based model to predict calprotectin depending on the patient group.

### 2.4. Patients

This study included two adult patients with a confirmed diagnosis of ulcerative colitis (UC). Patient 1 was a 41-year-old female of European descent. The disease was classified as left-sided colitis (E2 according to the Montreal classification) with a chronic relapsing course. At the time of the study, the disease activity index (DAI) score was 6. The disease duration was 11 years. There was no family history of UC. Her baseline therapy consisted of Guselkumab for 4 years and azathioprine for 7 years. Her medical history included more than three courses of corticosteroid treatment, with the last course administered 6 years prior to the study.

Patient 2 was a 36-year-old male of European descent. The disease was classified as extensive colitis (pancolitis, E2 according to the Montreal classification) with a chronic relapsing course. At the time of the study, the patient was in clinical remission with a DAI score of 7. Extraintestinal manifestations included primary sclerosing cholangitis, aphthous stomatitis. The disease duration was 7 years. There was no family history of UC. His baseline therapy consisted of Guselkumab for 4 years and azathioprine for 5 years. His medical history included corticosteroid treatment, with the last course administered 6 months prior to the study.

During a standard endoscopic procedure, biopsy samples were collected for analysis. From each patient, one sample was taken from the rectum and one from the ileum specifically for analysis by transmission electron microscopy (TEM). The present study reports exclusively on the TEM findings. Additional samples were collected for other analyses not reported here.

All procedures were reviewed and approved by the Ethics Committee of Novosibirsk State Medical University (Novosibirsk, Russia), protocol #164, 24 February 2025. All research was performed in accordance with the Declaration of Helsinki. Written informed consent was obtained from both participants involved in the study.

### 2.5. Animals

The experiments were performed in the Institute of Molecular and Cellular Biology (IMCB). All procedures were conducted under Russian legislation according to the standards of Good Laboratory Practice (directive # 267 from 19 June 2003 of the Ministry of Health of the Russian Federation), institutional Ethical committee guidelines and the European Convention for the protection of vertebrate animals. The experiments involving animal participants were reviewed and approved by the Ethics Committee on Animal and Human Research of the Institute of Molecular and Cellular Biology (Novosibirsk, Russia), protocol #02/21 dated 04 August 2021. All animals were tested for specific pathogens quarterly according to Federation of European laboratory animal science association’s (FELASA) recommendations (31). The study was conducted using *Muc2^−/−^* (*Muc2^tm1Avel^/Muc2^tm1Avel^*) mouse strain (32). Their wild-type littermates (*Muc2^+/+^* mice) were used as control.

Adult male mice were housed as same-sex siblings in open cages (with a dimension of 318 × 202 × 135 mm, #CP-3, 3 W, Russia) with birch sawdust as litter and paper cups as shelter. The housing conditions were as follows: 12 h/12 h light/dark photoperiod; and food (BioPro, Novosibirsk, Russia) and water were provided ad libitum.

Animals were euthanized by CO_2_ inhalation (30% of the chamber volume per minute). Descending colon samples were taken for RNA purification and frozen in liquid nitrogen until use, n=3 per group, and for electron microscopy, n=2 per group.

### 2.6. Transmission electron microscopy

Human colon and ileal samples were collected by a standard endoscopic procedure, n=2 per sample type. Samples were placed in a 2.5% glutaraldehyde solution in a 0.1 M sodium cacodylate buffer (pH 7.4) for 1 h at room temperature, washed and post-fixed in 1% osmium tetroxide with 0.8% potassium ferrocyanide for 1 h. Fixed samples were contrasted with 1% uranyl acetate in mQ water, dehydrated and embedded in epoxy resin (Epon 812). Mouse colonic samples were prepared similarly. Semithin cross-sections were prepared, stained with 1% methylene blue and analyzed with an Axioscope-4 microscope (Carl Zeiss). Ultrathin (70 nm) sections for TEM were obtained with a diamond knife (Diatome, Nidau, Switzerland) on a Leica EM UC7 ultramicrotome (Leica, Wetzlar, Austria) and then examined with a JEM1400 transmission electron microscope (JEOL, Tokyo, Japan). Section preparation and TEM was performed at the Center of Collective Use for Microscopic Analysis of Biological Objects (ICG SB RAS, Novosibirsk, Russia) (FWNR-2022-0015). Fixation, dehydration and embedding were performed at the Sector of Structural Cell Biology (ICG SB RAS).

### 2.7. RNA purification from murine colonic samples and trasncriptome analysis

The gene expression analysis was performed as follows. Total RNA was isolated from colon tissue samples using QIAzol lysis reagent (79306, Qiagen, Germany) and genomic DNA was removed by DNaseI (Roche, Germany) according to the manufacturers’ recommendations. RNA concentration was analyzed using a NanoDrop 2000 spectrophotometer (ThermoScientific, USA). Messenger RNA was purified from total RNA using poly-T oligo-attached magnetic beads. After fragmentation, the first strand cDNA was synthesized using random hexamer primers, followed by the second strand cDNA synthesis, adapter ligation and sequencing using an Illumina platform in a paired-end setting by Novogene (www.novogene.com, accessed on 01.08.2025) (33). Data and differential gene expression analysis were performed by Novogene company as follows: raw data were cleaned by removing reads containing adapters and low-quality reads. Paired-end clean reads were aligned to the reference genome using Hisat2 (version 2.0.5) (34). Differential expression analysis was performed using the DESeq2 R package (version 1.20.0) (35). The resulting P-values were adjusted using the Benjamini-Hochberg’s approach. Genes with an adjusted *p*-value < 0.05 were assigned as differentially expressed.

## 3. Results

### 3.1. Ultrastructural changes of the microvilli and the underlying actin filaments in UC patients resemble those found in model Muc2 mice with chronic colitis

Previous studies by our group and other authors have demonstrated that chronic colitis leads to ultrastructural alterations in the epithelium, particularly affecting cytoskeleton-associated components. To correlate qualitative changes in microvilli structure and actin filaments, we performed transmission electron microscopy (TEM) on colon tissue sections from *Muc2* knock-out (KO) mice ‒ a widely used model of chronic colitis and colorectal cancer ‒ and from patients diagnosed with ulcerative colitis. Inflamed tissues from both murine and human samples exhibited marked disruption of brush border ultrastructure (Figure 1). In healthy controls, microvilli demonstrated regular structure with parallel actin bundles, mostly enveloped by plasma membrane with only short uncovered rootlets comprising a clearly defined terminal web. In contrast, colonocytes affected by inflammation showed significant ultrastructural abnormalities including shortened and irregular microvilli with elongated rootlets (Figure 1). Core actin bundles of microvilli appeared to sink through the terminal web region in human samples similarly with our previous results on *Muc2* KO mice (6). Additionally, we observed organelles intruding the terminal web area (Figure 1, green asterisk). All these features indicate a persistent damage of the terminal web structure caused by chronic inflammation.

**Figure 1.**
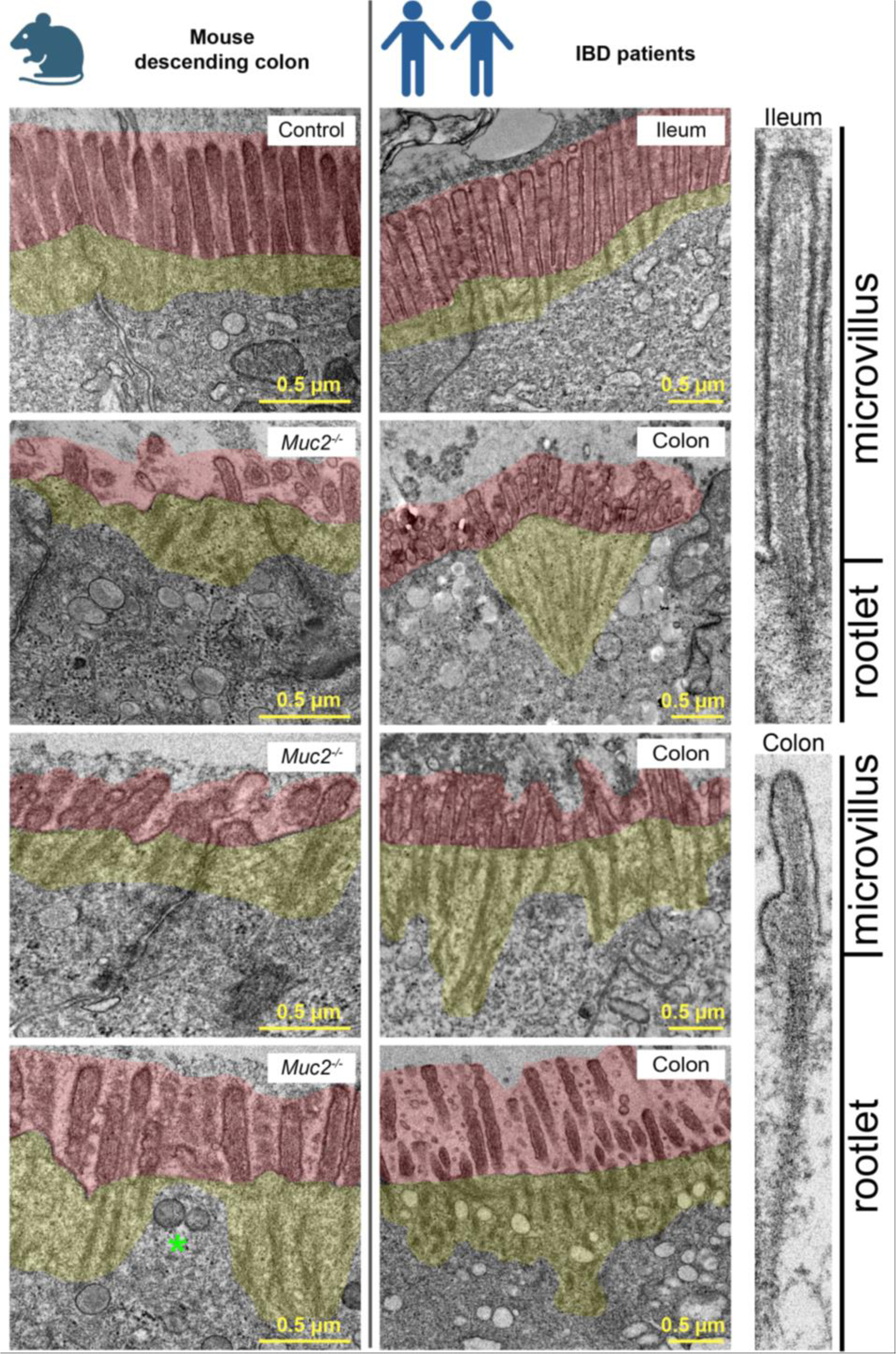
Ultrastructural analysis of the brush border in the descending colon of *Muc2*-deficient mice (n=2) and colonic biopsy from IBD patients (n=2). Healthy controls included the descending colon from wild-type mice and ileal biopsies from IBD patients, respectfully. Organelles intruding the terminal web area are marked by green asterisk. Microvilli and rootlets are highlighted with red and yellow, respectfully. Left panel shows murine colons, right panel depicts patients’ intestinal samples.

### 3.2. Bayesian network reveals key regulatory genes in patients with CD

The cohort used for Bayesian network (BN) composition consisted of 42 participants from Israel: 17 patients with diagnosed Crohn’s disease and 25 healthy controls with data available both for transcriptomic and metabolomic analyses, as well as clinical data. The data includes the gene expression measurements for 199,169 transcripts from the ileum, the abundances of 405 fecal metabolites, and 5 clinical characteristics, including the disease status, sex, age (patient’s age at the time of the study), blood C-reactive protein (CRP) and fecal calprotectin.

The subset of 42 participants used for further analysis had a diverse age distribution, with age windows of 15-29, 30-39, 40-49, 50-59, and 60-79 years. The mean CRP level in this cohort was 7.94 mg/L (SD = 10.35), with a median of 2.91 mg/L and a range from 0.28 to 38.97 mg/L. Fecal calprotectin levels showed substantial variation, with a mean of 448.49 µg/g (SD = 419.58), a median of 229.00 µg/g, and a range from 30.00 to 1000.00 µg/g.

Differential expression analysis revealed 1117 differentially expressed genes (DEGs) and 5 differentially abundant metabolites. Next, DEGs were split into 6 co-expression modules using WCGNA method and each module was correlated with metadata (Figure 2A). All modules correlated with calprotectin levels and CD status, while only blue, green, and turquoise modules correlated with CRP (Figure 2B). We then shortlisted candidate DEGs for the BN by identifying top 20 most influential hub genes in every module as evaluated by the combined influence score during WCGNA. Grey module only contained 24 genes, and was not used in further analysis. Immunoglobulin genes were excluded from the final list of genes, as they are not likely to be key regulatory components of the network, and are uneasy to consider as potential drug targets. Thus, only one gene from green module, *MZB1*, remained in the final list of genes. Thus, the resulting total number of 81 genes were used to build the BN (Figure 2C). Unfortunately, none of the metabolites formed significant connections within the network, so they were excluded from the network. The resulting BN exhibited a strong modular structure, with nodes grouping according to their original gene co-expression modules. This organization reflects the high correlation of expression profiles within each module. The network was organized hierarchically into four primary clusters. Each cluster was structured over 9-12 layers, governed by a master regulator gene at the apical node that subordinated the genes beneath it. Furthermore, the analysis revealed longer-range regulatory connections spanning genes from different modules. The network predicted four master regulator nodes: *PGRMC1* (brown module), *HEBP1* (blue module), *ANXA5* (turquoise module), and *PREX1* (yellow module) potentially controlling most of the network. The subnetwork derived from the turquoise module was the most interconnected, with the majority of its connections linking to the yellow module (Figure 2C).

**Figure 2.**
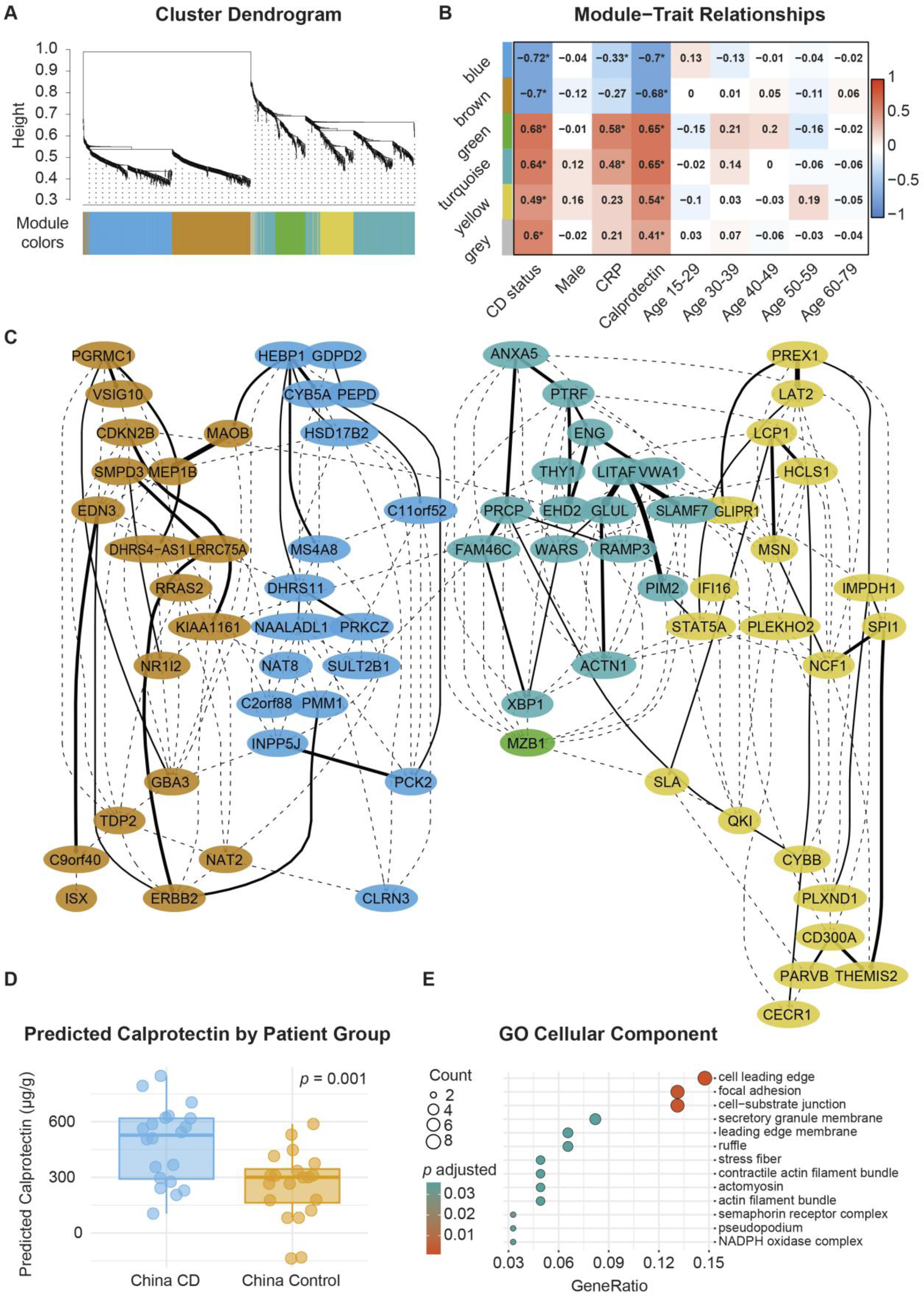
Construction and validation of a Bayesian Network (BN) from multiomic data in a Crohn’s disease cohort. (**A**) Cluster dendrogram showing gene modules identification by WGCNA. (**B**) Module-trait associations. Bar plot shows Pearson correlation coefficients between module eigengenes and clinical traits. \**p* < 0.05, \*\**p* < 0.01, \*\*\**p* < 0.001. (**C**) The final Bayesian network. The network was built using 81 genes and exhibits a hierarchical modular structure. Thicker lines define stronger connections, while dashed lines depict weaker interactions. (**D**) Validation of the BN model. Predicted calprotectin levels, generated by applying the BN model to transcriptomic data from an independent cohort (n=20 CD, n=20 controls), were significantly elevated in CD patients (*p*=0.001, Student’s t-test). (**E**) GO analysis of genes identified in the BN based on cellular component annotation.

In order to test the validity of the network, we incorporated calprotectin into the BN as a key clinical biomarker that reflects the severity of the intestinal inflammation. BN revealed one parent node of calprotectin ‒ *ISX*, that was further connected to the entire network by a gene with the unknown function *C9orf40* in the brown module. Subsequently, we constructed a predictive model based on this BN, aiming to evaluate its power in estimating calprotectin as a numerical factor. We see that in the Israeli patients (which was used as a training data set in BN-based model) transcriptomes explains approximately 63% of calprotectin variability that corresponds to Pearson’s correlation (between transcriptomic data and calprotectin) of r = 0.793 (p < 0.001). Next, we utilized the Chinese patient’s gene expression data as a test dataset to predict calprotectin using this newly built BN-based model. As there was no clinical calprotectin data for the Chinese sample provided in the original dataset, we divided the predicted numbers by patients group and tested statistical difference between them. Indeed, the BN-based model revealed significant upregulation of calprotectin in the patient’s group as compared to the control sample (n=20 in each group, China CD vs China control, t=3.55, *p* = 0.001, Student’s t-test, Figure 2D). This finding reveals that the BN-based predictive model can discriminate a major clinical parameter – calprotectin levels – between the two groups of patients, suggesting biological relevance of the BN and its gene connections.

### 3.3. Target Genes Shared in CD and a *Muc2* KO Colitis Model

Next, we analyzed gene ontology (GO) of the gene list revealed by the BN as nodes with strong connections. Interestingly, a majority of the identified GO terms linked these nodes to actin filaments and cytoskeleton-related structures (Figure 2E). In order to evaluate the relevance of these genes to microvilli disruption upon chronic intestinal inflammation, we used the *Muc2* KO mouse model of chronic colitis, which recapitulates key features such as microvilli disruption and cytoskeletal dysregulation. We have compiled a list of genes from the GO terms: cell leading edge, focal adhesion, cell-substrate junction, stress fiber, contractile actin filament bundle, actomyosin, and actin filament bundle found in CD patients (Table 1). Next, transcriptomic analysis of the intestinal gene expression profile in *Muc2* KO mice revealed the total list of DEGs in the mouse model of colitis. This allowed us to identify genes whose expression changes align with those of our target genes discovered through the BN. Mouse gene expression data of the distal colonic tissue identified over a thousand DEGs between *Muc2* KO mice and wild type littermates (Figure 3A, 3B). Mouse gene expression data was directly used for GO analysis as a full list of DEGs with no further pre-selection, which revealed no cytoskeleton-specific terms (Figure 3C). Then, we examined the expression of the mouse orthologs of the pre-selected human genes (Table 1). Notably, we found only one gene, *PREX1*, that changed expression in both CD patients and *Muc2* KO mouse model, so that the direction of change was consistent. Given that *PREX1* is a key player in the Rho signaling pathway ‒ which modulates both inflammation and actin dynamics ‒ and a top regulatory node in our BN subnetwork, it can be considered a primary target for controlling microvilli structure and epithelial integrity in chronic colitis.

**Figure 3.**
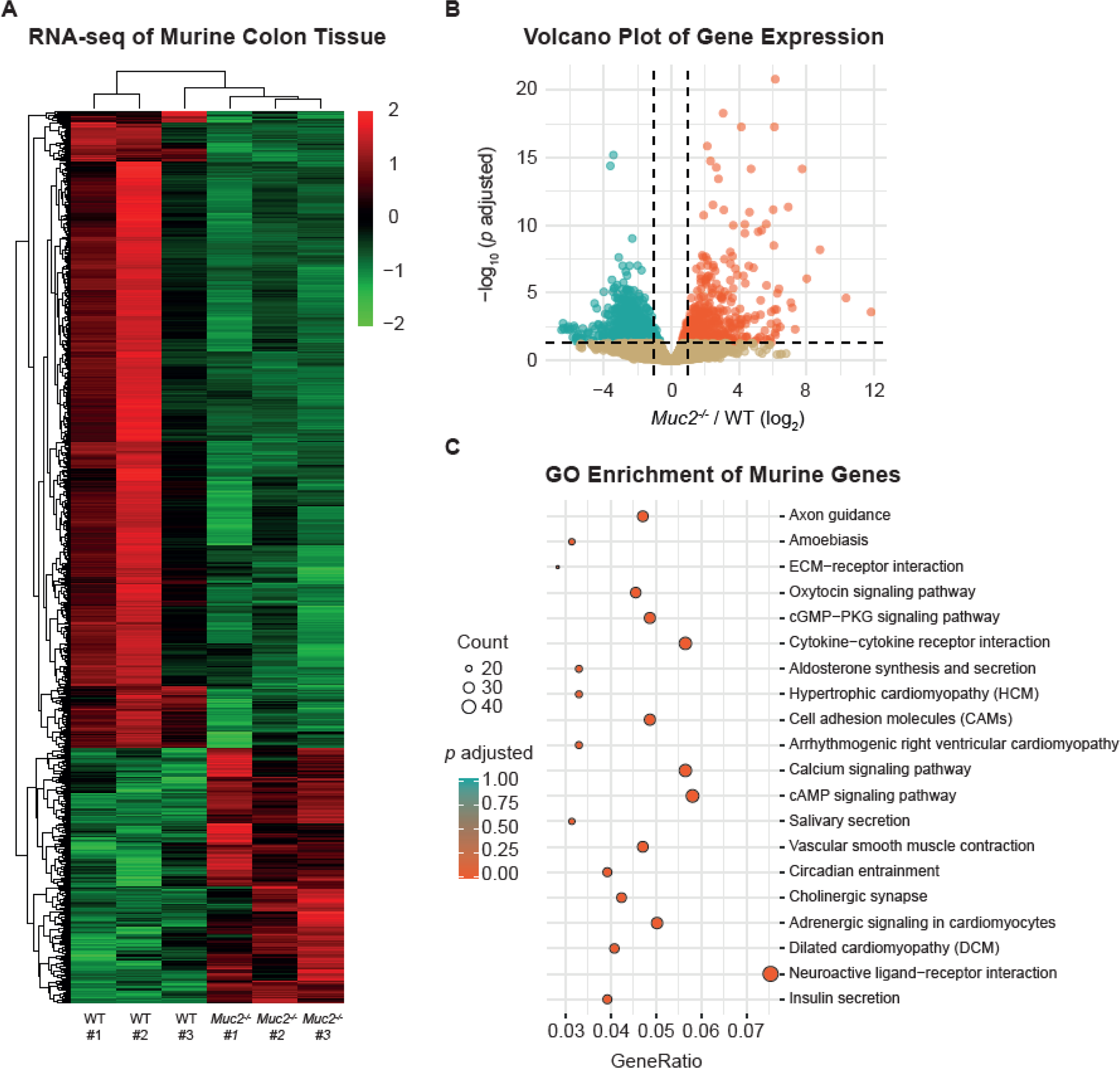
Identification of DEGs in the distal colon *Muc2* KO mouse model of chronic colitis. (**A**) Clustered heatmap showing DEGs in individual biological replicates. (**B**) Volcano plot showing averaged DEGs per group (n=3 per group, the difference was calculated with DESeq2 using a negative binomial generalized linear model (35), and corrected by FDR). (C) GO enrichment analysis for the entire list of DEGs with significant difference in expression.

**Table 1.**
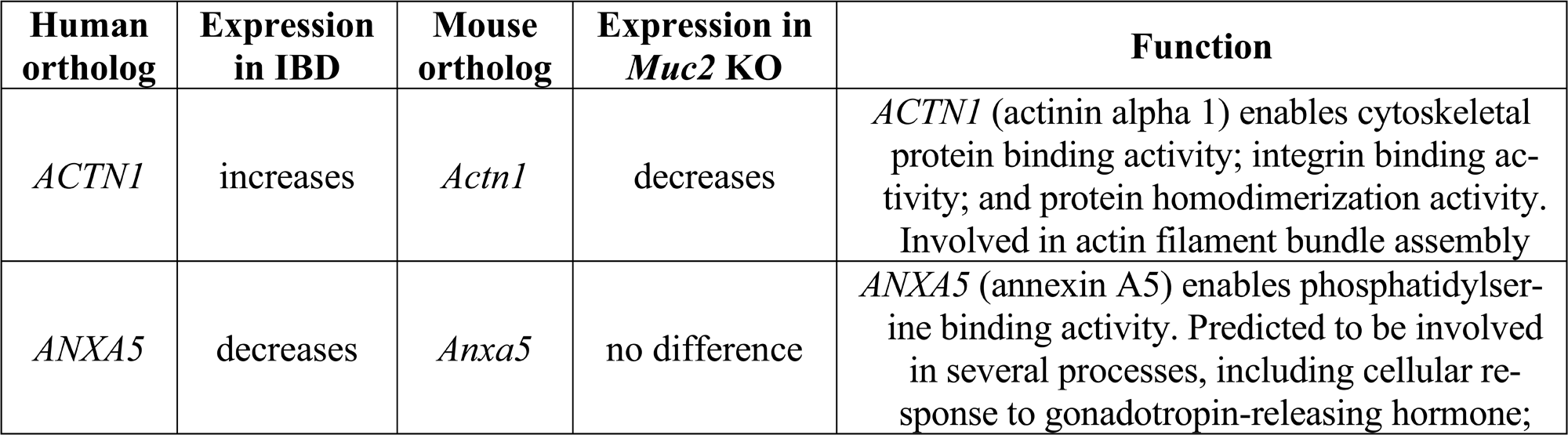
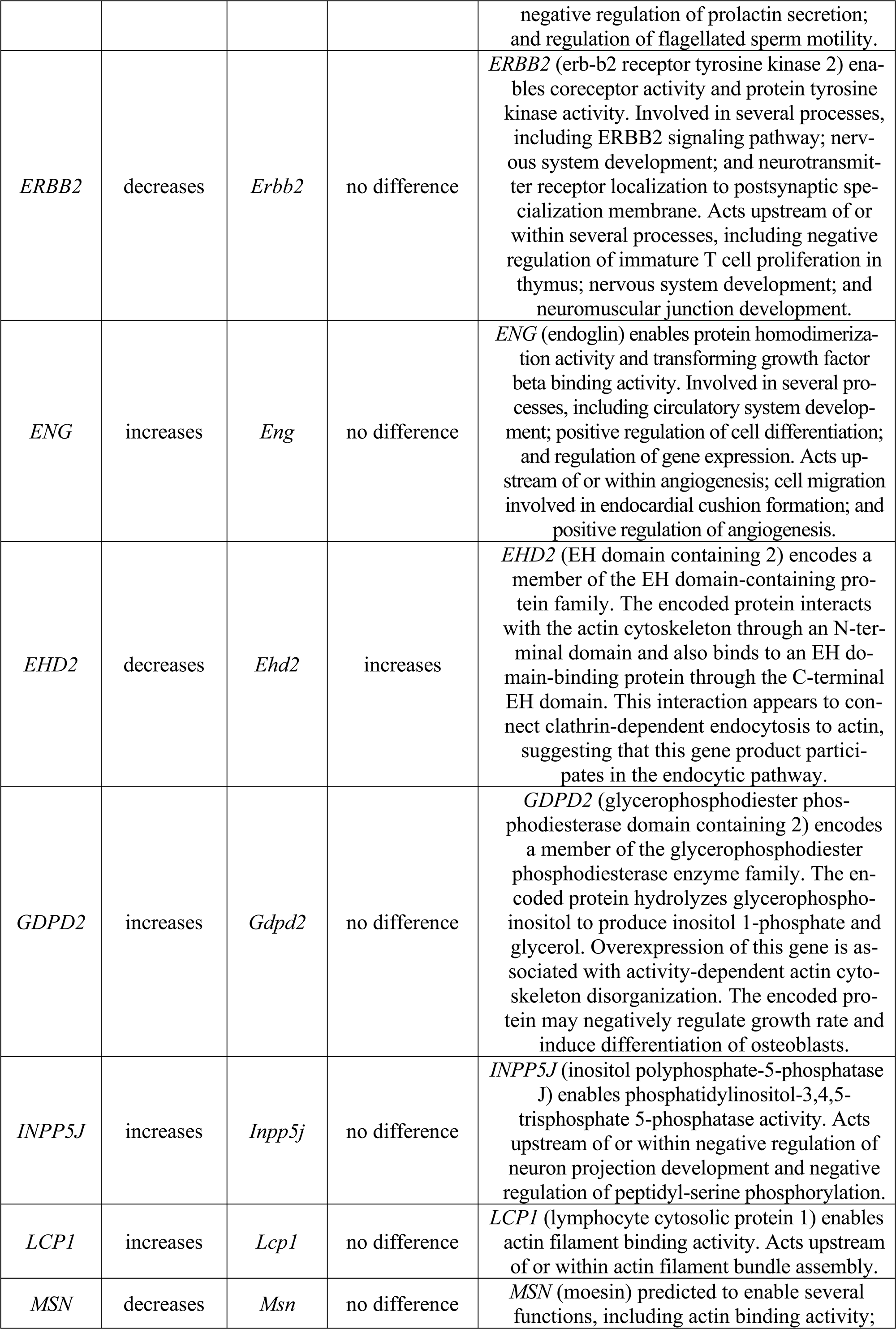
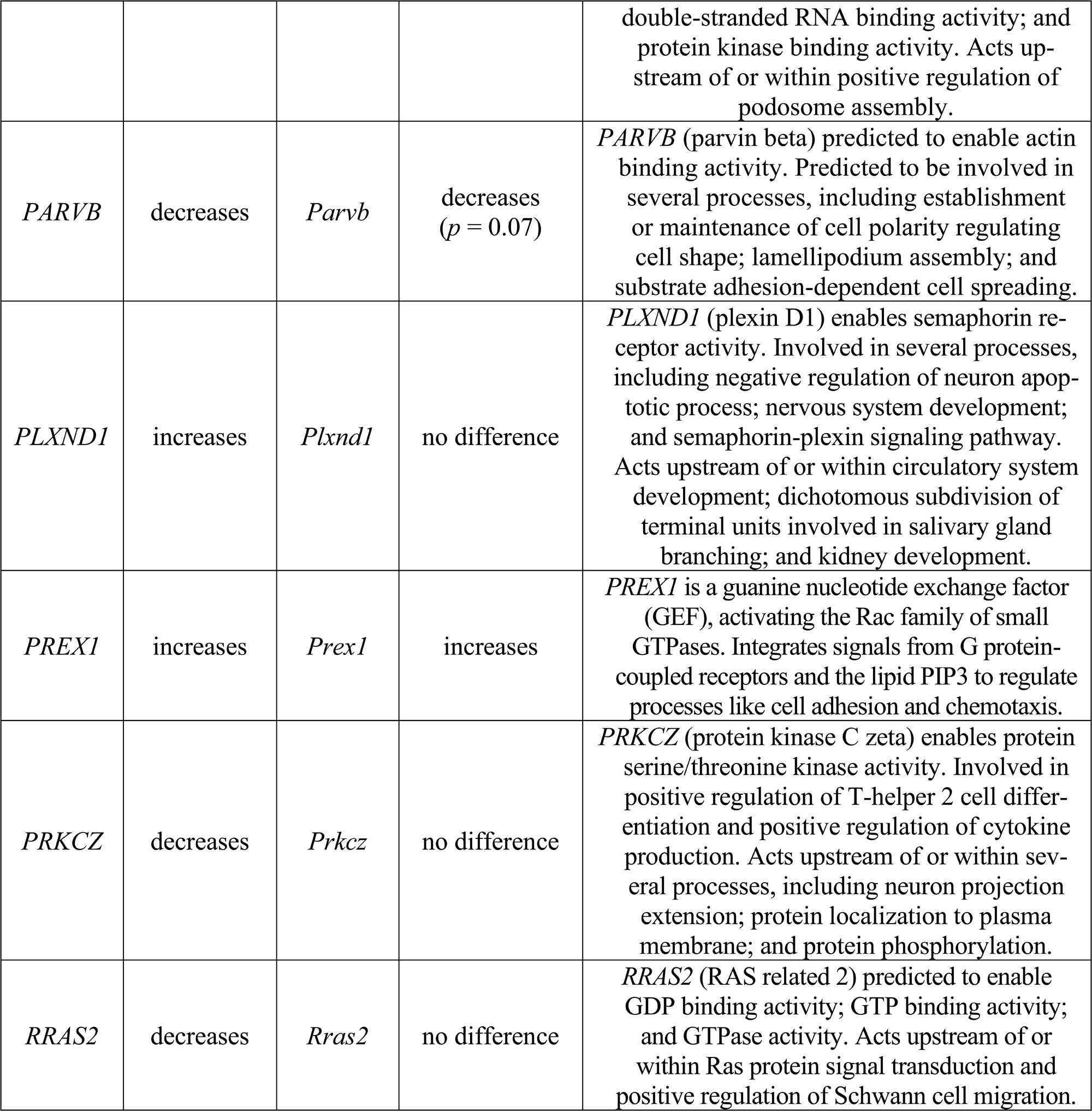
Expression analysis of human gene orthologues in *Muc2* KO mice.

## 4. Discussion

Brush border disorganization is associated with a range of gastrointestinal pathologies, including infectious and genetic inflammatory conditions (36). Ultrastructural analysis of intestinal biopsies from patients with celiac disease demonstrated severe damage of enterocytes, accompanied by destruction of microvilli, rootlets, and the terminal web (37,38). Microvillar fusion, as well as decreased microvillar length and density have been discovered in pediatric IBD samples (39). VanDussen et. al. reported that significant loss of the microvilli rootlets in CD patients was associated with downregulation of several key genes involved in brush border organization, namely *PLS1*, *MYO5B*, *STX3*, and *USH1C* (8). Experimental models of intestinal inflammation, such as DSS-induced colitis, also replicate these morphological defects, demonstrating substantial microvillar disruption and loss of epithelial integrity (40,41). These data suggest that an inflammatory environment induces significant changes in brush border architecture causing disruption of the intestinal absorptive surface.

In this study, we describe a remarkable ultrastructural phenotype of intestinal epithelium in both *Muc2* KO IBD model and human IBD patients. Transmission electron microscopy revealed microvillar shortening accompanied by elongation of the rootlets and widening of the terminal web area. A comparable terminal web widening has been previously reported in duodenal epithelium of *Cobl* KO mice (42). Dysfunction of the actin nucleator Cordon-bleu (*Cobl*) leads to the loss of fine filaments (likely, loose F-actin network) that connect the cell cortex, the rootlets and the basal intermediate filaments of the terminal web (42). Syndapin-2, also known as Pacsin2, which is involved in membrane remodeling, has been shown to recruit Cobl to the apical domain of the terminal web (43). Pacsin2 also interacts with N-WASP, an activator of the Arp2/3 complex, implicating its involvement in the actin polymerization process (44). Loss of Pacsin2 in murine duodenal epithelium leads to significant reduction in the ratio of microvillus length to total actin bundle length, resembling results shown in *Cobl* KO or *Muc2* KO mice (45). Additionally, inducible deletion of the small GTPase Rab11a in neonatal intestinal epithelial cells results in a similar ultrastructural change, consistent with its critical role in trafficking brush border proteins, including syndapin-2 and villin (46).

Our transcriptomic analysis of *Muc2* KO mice and IBD patient transcriptomic profiles identified *PREX1* as a candidate gene associated with brush border remodeling. The mouse ortholog *Prex1* was found to be upregulated in the murine IBD model, consistent with observations in human samples. PREX1 is a guanine nucleotide exchange factor for Rac GTP-binding proteins which belong to the Rho family of small GTPases involved in actin dynamics. PREX1 activity is regulated by several mediators including PIP3 and Gβγ, the sphingosine-1-phosphate receptor S1P1, cAMP-dependent protein kinase PKA, the protein phosphatase 1α, and mammalian target of rapamycin (mTOR), which are known to be activated during inflammation (47,48). Rac GTPases encompass 4 members: RAC1 which is ubiquitous, RAC2 which is mainly expressed in hematopoietic cells, RAC3 which is expressed in brain and testis, and RHOG which is present in fibroblasts, leukocytes, neuronal, and endothelial cells (49). It’s been demonstrated that PREX1 binds and activates RAC1 (47,48). In primary human T-cells, active RAC1 induces dephosphorylation of the ezrin/radixin/moesin family of actin regulators, leading to microvilli disassembling (50). *Rac1* is upregulated in DSS-induced colitis, whereas pharmacological inhibition of its activity ameliorates clinical symptoms and epithelial damage (51). PACSIN2-mediated endocytosis is negatively regulated by RAC1 signaling (52). Presumably, elevated PREX1 levels in IBD could lead to RAC1 overactivation and suppression of PACSIN2 activity, ultimately destabilizing cortical actin and causing a shortening of microvilli in epithelial cells (45). From the other perspective, downregulation of Rac1 results in dramatic cytoskeleton rearrangement, impaired cell extrusion and intestinal permeability (4) suggesting the importance of maintaining the balance of RAC1 activity.

*PREX1* also has a role in tumorigenesis and cancer metastasis. Elevated *PREX1* expression has been reported in cancers, including melanoma and breast, prostate, and colon cancer (53). Downregulation of *PREX1* expression suppresses breast cancer cell growth (48). The *RAC1* high-risk allele has been associated with an increased chance of colorectal cancer developing in IBD patients (54) which can be explained by its role in the Wnt/β-catenin (49). RAC1 and PREX1 contribute to epithelial-mesenchymal transition in gastric cancer (55).

In addition to *PREX1*, our analysis identified actin-binding proteins which potentially can have an effect on microvilli ultrastructure, namely *ACTN1* and *EHD2*. Surprisingly, both genes were downregulated in the *Muc2* KO mice but upregulated in human IBD biopsies. Such discrepancies may arise from differences in the inflammatory environment between murine IBD model and human disease with distinct mechanisms underlying disease onset and progression. However, the involvement of these genes in both *Muc2* KO colitis and human IBD suggests that an alteration in the expression of actin-regulating proteins can substantially disrupt the cytoskeleton organization and cell morphology.

*ACTN1* encodes α-actinin-1, an F-actin crosslinking protein that maintains cortical cytoskeleton organization, and mediates interactions of cytoskeletal regulatory proteins (e.g., capping protein, vinculin, zyxin, integrin, cell surface receptors, etc.) with the cell membrane (56). Actinin 1 is localized in the vicinity of cell junctions (57,58). Several earlier studies have reported that α-actinin-1 is distributed between microvilli rootlets (59,60), yet more recent evidence on its localization in the terminal web is lacking. Given the presence of Cobl in this compartment, α-actinin-1 may participate in the loose actin mesh interconnecting microvillar rootlets, but further studies revealing the terminal web structure are required. Studies have shown that *ACTN1* expression is altered in IBD patients (61), its upregulation disrupts E-cadherin-based adhesions compromising epithelial integrity (62).

EHD2 attaches F-actin to the plasma membrane acting as a negative regulator of clathrin-mediated endocytosis (63). This protein interacts with PACSIN2 and coordinates membrane remodeling and vesicle scission (64,65). Loss of EHD2 causes the detachment of caveolae from the plasma membrane (66). Similarly with vesicle-scissoring PACSIN2, alterations in EHD2 expression may lead to higher frequency of membrane invaginations in the intermicrovillar region in brush border in IBD patients (45). No consistency in *EDH1* expression and its mouse orthologue was found in CD patients and the mouse model.

One of the top regulator genes revealed by the BN is *ANXA5*. It is a calcium-dependent phospholipid-binding protein that orchestrates key cellular processes, most notably through its direct regulation of the actin cytoskeleton, which is fundamental to maintaining gut epithelial integrity (67). This protein binds to actin filaments in a calcium-sensitive manner, thereby stabilizing the cortical actin ring, promoting membrane resealing during injury, and facilitating restitution of the epithelial barrier (68). By anchoring the cytoskeleton to phospholipid membranes, ANXA5 coordinates actomyosin contractility during cell migration, a process critical for wound healing in the gut mucosa (69). ANXA5 alleviates experimental colitis by targeting phosphatidylserine (PS)-exposed colonic capillaries, inhibiting endothelial cell activation and TLR4-mediated inflammation through PS-dependent endocytosis, highlighting PS as a target for designed drug delivery in IBD (70). Together with published results, ANXA5 can be considered a pivotal regulator of cytoskeletal-driven intestinal integrity and a potential biomarker for IBD severity and progression. However, mouse transcriptomic data suggest that *ANXA5* orthologue’s expression is not affected in the *Muc2* KO model of colitis, which makes it an unlikely regulator of microvilli integrity.

Another critical gene is *ISX*, identified as a key effector hub in our analysis. ISX is an intestinal transcription factor that acts as a molecular nexus, integrating dietary signals ‒ particularly the availability of β-carotene (pro-vitamin A) ‒ with mucosal immune responses (71,72). This protein directly regulates the expression of the enzyme BCO1, which converts β-carotene into retinal (vitamin A), and controls the subsequent production of retinoic acid in the gut (71,72). Retinoic acid is a critical signaling molecule that promotes the homing of immune cells to the gut mucosa and is essential for the differentiation and maintenance of regulatory T cells (Tregs) and the generation of IgA-secreting plasma cells (73,74). Moreover, genetic polymorphisms in the *ISX* have been associated with inflammatory bowel disease in a genome-wide association study (75). Thus, our model reveals *ISX* as a potential key effector of mucosal immunity and a potential therapeutic target in IBD.

## 5. Conclusions

Our BN analysis identified *PREX1*, *ACTN1*, *EHD2*, and *ANXA5* as candidate regulators of brush border organization in CD patients’ transcriptomic data. Only *PREX1* dysregulation coincides with ultrastructural disruption of the microvilli-terminal web complex in both *Muc2* KO colitis and IBD patients. By linking transcriptional changes to defined structural defects in human patients and model mice, our findings identify PREX1 as one of the new potential regulators of epithelial damage and possible drug targets for the treatment of IBD.

## Author Contributions

Conceptualization, E.K. and M.S.; formal analysis, E.K. and S.M.; investigation, S.M., J.K., K.M., J.P., and M.S.; data curation, M.O. and M.S.; writing ‒ original draft preparation, S.M. and E.K; writing ‒ review and editing, S.M., J.K., K.M., J.P., M.S., M.O., and E.K.; visualization, S.M., J.P., and E.K.; supervision, M.S., M.O., and E.K.; project administration, E.K.; funding acquisition, E.K. All authors have read and agreed to the published version of the manuscript.

## Informed Consent and Ethical Statement

The study involving human participants was reviewed and approved by the Ethics Committee of Novosibirsk State Medical University (Novosibirsk, Russia), protocol #164, 24 February 2025. All research was performed in accordance with the Declaration of Helsinki. Written informed consent was obtained from both participants involved in the study. The experiments involving animal participants were reviewed and approved by the Ethics Committee on Animal and Human Research of the Institute of Molecular and Cellular Biology (Novosibirsk, Russia), protocol #02/21 dated 04 August 2021.

## Data Availability Statement

RNASeq Israel and China datasets used in this study have been downloaded from the Gene Expression Omnibus database under accession code: GSE199906 and GSE233900, respectively. These data were derived from the original study by Braun et. al., 2024 (23). Mouse transcriptomic data obtained in this study is available to the reviewers by request and will be deposited to the corresponding database upon publication.

## Acknowledgments

TEM analysis was performed at the Center of Collective Use for Microscopic Analysis of Biological Objects (ICG SB RAS, Novosibirsk, Russia, FWNR-2022-0019). The authors employed a Large Language Model (DeepSeek-v3, https://www.deepseek.com/) for proofreading and editing the English language in this manuscript.

## Abbreviations

The following abbreviations are used in this manuscript:

IBD: Inflammatory bowel disease
CD: Crohn’s disease
UC: Ulcerative colitis
DEGs: Differentially expressed genes
WGCNA: Weighted correlation network analysis
BN: Bayesian network
CRP: C-reactive protein
PS: Phosphatidylserine

